# Oncogenic chromodomain mutations allosterically impair TIP60 acetyltransfearse function preventing activation of DNA repair genes under genotoxic stress

**DOI:** 10.64898/2026.02.23.707364

**Authors:** Himanshu Gupta, Aarushi Bansal, Ashish Gupta

## Abstract

TIP60 is a tumor suppressor with histone acetyltransferase activity that regulates chromatin accessibility in diverse processes including DNA repair, apoptosis, mitosis, transcription, and autophagy. Structurally TIP60 contains a C-terminal MYST domain that mediates its HAT activity, while the N-terminal chromodomain are conserved modules that facilitate its association with chromatin by recognizing histone modifications. Mutations within the chromodomain have been implicated in various cancers, yet their functional consequences remain poorly understood, particularly with respect to TIP60’s role in maintaining genomic integrity. Here, we uncover a novel allosteric mechanism whereby cancer-associated chromodomain mutations impair TIP60’s catalytic activity without disrupting its chromatin binding, underscoring critical interdomain communication between the chromodomain and the MYST domain. Through structural modelling and molecular dynamics simulations, we identified two missense mutations (R53H and R62W) in TIP60’s chromodomain that not only altered TIP60’s conformation but also destabilized its trimeric assembly, thereby impairing acetyl-CoA docking. Importantly, we found that TIP60 engages acetyl-CoA exclusively in its trimeric state, and the R62W mutation perturbs the trimeric interface, thereby altering the docking sites for acetyl-CoA. Consistent with these structural changes, biochemical assays revealed that chromodomain mutant TIP60 variants, while retaining chromatin loading, exhibited markedly reduced autoacetylation and histone acetyltransferase activity. Moreover, these mutants failed to activate the *p21* gene in response to DNA damage, thereby predisposing the genome to the accumulation of mutations and leaving cells unable to arrest the cell cycle for repair of genomic lesions. Together, our findings establish that distal chromodomain mutations allosterically destabilize TIP60 oligomerization, impair acetyl-CoA utilization, and compromise DNA damage responses. The mechanism establishes a link between chromodomain mutations and genomic instability, shedding light on how reader domain alterations may underlie cancer progression.

## INTRODUCTION

Tat-interactive protein 60 (TIP60; KAT5), a MYST family histone acetyltransferase is a critical regulator of chromatin architecture and genome stability (Sapountzi et al., 2006). Originally identified as a cofactor for HIV-1 Tat-mediated transcription, TIP60 is now recognized for its role in transcriptional regulation, DNA damage repair, apoptosis, cell cycle progression, stem cell identity, and lineage differentiation (Acharya et al., 2017; Hlubek et al., 2001; Ikura et al., 2000; Kamine et al., 1996; Li et al., 2012; Numata et al., 2020). As a lysine acetyltransferase, TIP60 mediates the transfer of acetyl groups from acetyl-CoA to histone lysine residues, leading to chromatin remodeling and coordinated transcriptional regulation. (Sapountzi et al., 2006). Beyond histones, TIP60 also acetylates key non-histone substrates involved in the DNA damage response, including the tumour suppressor p53 and the ATM kinase, a DNA damage sensor (Gupta & Gupta, 2025; Sun et al., 2005; Tang et al., 2006). These modifications facilitate the coordination of repair pathways and influence cell fate decisions such as apoptosis and cell cycle arrest (Gupta & Gupta, 2025; Li et al., 2012; Squatrito et al., 2006; Sun et al., 2010).

The functional versatility of TIP60 arises from the interplay of its distinct structural modules. Structurally, TIP60 comprises three major domains, that includes an N-terminal chromodomain (CD), an intrinsically disordered region (IDR), and a C-terminal MYST acetyltransferase domain that forms its catalytic core (Gupta & Gupta, 2025; Sapountzi et al., 2006). The MYST domain harbors the enzymatic machinery, including a zinc finger motif essential for substrate recognition and acetyltransferase activity (Mir et al., 2021; Sapountzi et al., 2006; Tam et al., 2017). Though the chromodomain and the MYST domains are spatially distinct, they are connected by flexible intrinsically disordered region (IDR). TIP60’s IDR contributes to phase separation, supporting the assembly of dynamic nuclear compartments or condensates that concentrates regulatory factors (Dubey et al., 2024.). Chromodomain functions primarily as methyl-lysine readers and guide diverse protein complexes to their sites of action (Eissenberg, 2012). For instance, HP1 chromodomain bind H3K9me3 to enforce heterochromatin and silence repetitive elements (Eissenberg, 2001; Zeng et al., 2010). Within Polycomb CBX proteins such as CBX7, the chromodmain recognize H3K27me3 to maintain transcriptional repression during developmental programming (Ma et al., 2014). Likewise, TIP60’s chromodomain mediates its binding to methylated lysine residues on histone tails, thereby directing TIP60 to specific chromatin loci to ensure that its catalytic activity is deployed precisely where transcriptional regulation or DNA damage response is required (Kim et al., 2015; Sun et al., 2009).

Large-scale sequencing efforts is increasingly uncovering missense mutations in the genome of cancer patients, and driving efforts to clarify their functional roles in tumor development and progression (Cerami et al., 2012; Gao et al., 2013; Sharma et al., 2025; Zhao et al., 2018). Among these, mutations within chromatin-associated domains are of particular interest, due to the prevailing assumption that CD mutations primarily weaken or disrupt chromatin association, thereby impairing downstream transcriptional programs and contribute to malignant transformation. For example, mutations in the CHD4 chromodomain have been identified in endometrial and colorectal cancers, where they disrupt nucleosome mobilization and DNA double-strand break repair (Li et al., 2018; Lin et al., 2019). Likewise, pathogenic missense mutations in the CBX7 chromodomain can weaken Polycomb-mediated repression, leading to inappropriate gene reactivation and oncogenic transformation (Clermont et al., 2014; Ren et al., 2015). Although the full consequences of these alterations remain under investigation, it is critical to determine whether mutations within the TIP60 chromodomain affect its structural and functional dynamics including its localization, enzymatic activity, and regulation of pathways such as the DNA damage response.

In this study, we investigated the impact of cancer-associated chromodomain mutations in TIP60. Using long-timescale molecular dynamics simulations, structural modelling, and biochemical assays, we identified two destabilizing missense mutations (R53H and R62W) that do not impair chromatin binding but instead destabilize TIP60’s conformational stability and disrupt TIP60’s trimeric assembly and acetyl-CoA docking. These defects led to reduced autoacetylation, diminished HAT activity, and failure to activate downstream DNA damage response genes such as p21. These findings indicate that chromodomains are not merely chromatin-binding modules but also exert allosteric control over TIP60’s catalytic function by facilitating the formation of its trimeric oligomers and thus revealing a previously unrecognized mechanism of interdomain communication between the CD and MYST domain. By delineating this unexpected allosteric mechanism, our study provides new insight into how cancer-associated mutations in TIP60 contribute to oncogenesis and highlights the broader principle of functional interdependence among distinct domains of chromatin regulators.

## Material and methods

### Identification and *in silico* assessment of deleterious and destabilizing mutations

Data curation and prediction of deleterious and destabilizing mutations followed the protocol described by Gupta et al. (Gupta et al., 2025). KAT5 gene mutation data were retrieved from cBioPortal (http://www.cbioportal.org) using its rapid search feature, focusing on alterations in TIP60 across various cancers; this identified six missense mutations in its chromodomain. COSMIC database data yielded 22 missense mutations. Six overlapping mutations common to both databases were selected for prediction analysis. PredictSNP assessed deleterious effects by integrating MAPP, PhD-SNP, PolyPhen-1, PolyPhen-2, SIFT, and SNAP tools. Stability predictions utilized I-Mutant 2.0, SAAFEC-SEQ, and INPS, calculating ΔΔG values where negative scores indicate destabilization and positive scores indicate stabilization.

### Structure preparation and refinement

The TIP60 N-terminal chromodomain structure (residues 1-78) was modeled using TTARoseFold on the Robetta web server, generating five models; the highest-quality model was refined with GalaxyRefine from the Galaxy WEB server. Model quality was validated via SAVES v5.0, incorporating Verify-3D (requiring 3D-1D scores ≥1.0 per residue for accuracy), ERRAT (assessing atomic interactions to distinguish correct from incorrect regions), and PROCHECK (evaluating overall and residue-specific stereochemical geometry). Mutations were introduced using PyMOL’s amino acid substitution plugin, with PyMOL also facilitating structure visualization and superimposition.

### Molecular dynamic simulation

A Molecular Dynamics (MD) simulation was conducted using Desmond by D.E. Shaw to fully understand the effects of mutations on the conformational stability of the TIP60 N-terminal chromodomain (1–78) wild-type and point mutants (R53H or R62W). A TIP3P water model was used to solvate the system for molecular dynamics simulations. Periodic boundary conditions were applied using a cubic simulation box with a 10 Å buffer distance around each system. Chloride counterions were added to ensure overall charge neutrality. Desmond’s standard relaxation procedure was employed to optimize and minimize the systems. The equilibrated systems were simulated utilizing the NPT ensemble at 300 K and 1.01325 bar, with a time step of 2 fs. Each molecular dynamics simulation for an apo protein was conducted for 500 nanoseconds, with energy and trajectory data recorded every 1.2 picoseconds and 100 picoseconds, respectively.

### Cell culture, transfection, and live cell imaging

Cos-1 cells (monkey kidney fibroblasts) and Huh7 cells (human hepatoma cells) were cultivated in DMEM enriched with 10% fetal bovine serum and 0.5% penicillin/streptomycin antibiotic solution. Cells were sub-cultured after they reached 80% confluency utilizing 0.5% trypsin EDTA and were sustained at 37°C in humid conditions with 5% CO supplementation. The Cos-1/Huh7 cells were transfected with desired plasmids utilizing Lipofectamine 2000 reagent (Invitrogen) according to the manufacturer’s instructions. Following a five-hour period, the medium was replaced with DMEM supplemented with 5% FBS and incubated for 24 hours. A fluorescence microscope (Nikon Ti Eclipse) was employed to ascertain the localization of fluorescent-tagged proteins in transfected cells. DAPI (4′,6-diamidino-2-phenylindole) was used to visualize the nucleus.

### Cloning

TIP60 mutants were created using overlapping PCR to change the amino acids arginine at position 53 and arginine at position 62 to histidine and tryptophan, respectively. The TIP60 (mutant) open reading frame was amplified utilizing mutation-specific primers in PCR-1 and PCR-2 reactions. The outputs of PCR-1 and PCR-2 served as the template for PCR-3, using TIP60 wild-type primers, which further amplified the TIP60 (mutant) open reading frame (ORF). The amplified ORF was purified utilizing a PCR cleanup kit (Wizard SV gel and PCR clean-up kit). The purified amplicon was digested with KpnI and BamHI enzymes using standard restriction digestion protocol prior to ligation into the digested pDsRed vector utilizing T4 DNA ligase. Subsequently, DH5 alpha competent cells were transformed with the ligation mixture, and the bacterial cells were then plated on the LB agar plate and kept at 37°C for 12-14 hours. The obtained colonies were then screened for positive clones using a double digestion. Once positive clones were obtained, generation of mutation was confirmed by sequencing. To construct pET28a-TIP60 (R53H) or pET28a-TIP60 (R62W) clone in the pET28a vector, PCR amplification was conducted to obtain the full-length ORF of TIP60 using specific primers and RFP-TIP60 (R53H) or RFP-TIP60 (R62W) as a template, subsequently followed by restriction digestion of the amplified PCR product and vector utilizing BamHI and EcoRI restriction enzymes. Ligation was executed utilizing T4 DNA ligase to produce the construct. Generated clones were validated through restriction enzymes digestion. The pDSRed-TIP60 (wild-type) and pET28a-TIP60 (wild-type) clones were previously generated in the laboratory. The primers utilized in the study are listed in the supplementary data (Supplementary Table 1).

### Western blotting

Mammalian cell lysate or purified protein samples were subjected to heat denaturation using 2X Laemmli sample buffer prior to separation via SDS-PAGE for Western blot analysis. Proteins on the gel were subsequently transferred to a methanol-activated PVDF membrane utilizing a semi-dry transfer apparatus. The PVDF membrane was subsequently blocked with 5% non-fat dried milk for one hour on a low-speed rocker at ambient temperature. The membrane was washed with 1X PBST three times for five minutes each. The membrane was subsequently incubated with the primary antibody with gentle agitation overnight at 4°C. The blot was subsequently washed three times with 1X PBST and incubated with HRP-conjugated secondary antibody for one hour at room temperature, followed by washing with 1X PBST. Ultimately, signals were developed utilizing the ECL reagent and were captured and analysed using a Chemi-Doc system. All the antibodies used are mentioned in Supplementary Table 2.

### Recombinant protein purification

Protein purification was conducted as described by Dubey et al. (Dubey et al., 2022). In short, to purify recombinant proteins, BL21 (DE3) cells were transformed with pET28a-TIP60 (wild-type), pET28a-TIP60 (R53H), or pET28a-TIP60 (R62W) plasmid constructs via a standard heat shock protocol. The isolated transformed bacterial colony was subsequently selected from the Luria Bertani (LB) agar plate and inoculated into LB broth containing kanamycin, incubated at 37°C at 200 rpm for 12–14 hours, constituting the primary culture. After inoculating the secondary culture with 1% of the primary culture, the cells were cultivated until the optical density reached 0.6. Recombinant protein induction was then initiated using 0.5 mM IPTG for 16 hours at 16°C. Subsequent to induction, cells were pelleted via centrifugation at 5000 rpm for 10 minutes at 4°C. Cells that were harvested were then lysed in a lysis buffer composed of 1X PBS, 2 mM EDTA, 5 mM DTT, 0.5 mM PMSF, 0.1% Triton X-100, 10% glycerol, 10 mM imidazole, and 100 μg lysozyme, followed by sonication using a probe sonicator. Centrifugation was subsequently conducted to obtain a clear supernatant. The Ni-NTA beads were equilibrated with lysis buffer, subsequently added to the clear supernatant, and incubated for one hour with slow rotation at 4°C to facilitate the binding of the His-tagged protein. The Ni-NTA beads were retrieved by centrifuging the supernatant at 2000 rpm for 5 minutes and subsequently washing the beads with wash buffer (1X PBS, 0.1 mM PMSF, 20 mM imidazole). The protein adhered to the beads was subsequently eluted using an elution buffer composed of 50 mM Tris at pH 8.0, 10% glycerol, 150 mM NaCl, and 500 mM imidazole. The eluted protein underwent dialysis for 4 hours in HAT buffer at 4°C, after which the dialyzed proteins were employed for *in vitro* experiments.

### *In vitro* HAT and autoacetylation assay

The assays were conducted in accordance with the previously established protocol (Bakshi et al., 2017). In short, for the HAT assay, purified recombinant proteins were incubated with acetyl CoA and 1 microgram of H4 peptide in HAT buffer for 1 hour at 30°C using a dry bath. For the autoacetylation assay, purified recombinant proteins were incubated with acetyl CoA and reaction buffer for 1 hour at 30°C. The reactions were halted by the addition of 2X Laemmli buffer and subsequently subjected to heat denaturation at 95°C for 10 minutes. The samples were analyzed using SDS-PAGE, followed by Western blotting with appropriate antibodies.

### Subcellular fractionation

The subcellular fractions were prepared as previously described (Gupta et al., 2025). Briefly, Cos-1 cells were transfected with RFP-TIP60 (wild-type), RFP-TIP60 (R53H), or RFP-TIP60 (R62W) constructs using Lipofectamine 2000. After 24 hours of media replacement, cells were trypsinized and harvested for subcellular fractionation. To obtain the soluble fraction, comprising cytoplasmic and soluble nuclear components, cells were lysed in soluble lysis buffer (10 mM HEPES, pH 7.4; 10 mM KCl; 0.05% NP-40; 0.2 mM MgCl ; 1% Triton X-100; 100 mM NaCl; 1× PIC) and incubated on ice for 20 minutes. Subsequently, the cell lysate was centrifuged at 1300g for 5 minutes at 4°C, and the supernatant, containing the soluble fraction, was collected. The cell pellet was then rinsed twice with a soluble lysis buffer. To isolate the chromatin-bound fraction, the residual cell pellet was lysed in chromatin lysis buffer (50 mM Tris pH-8.0, 400 mM NaCl, 10 mM EDTA, 0.5% SDS, 1X protease inhibitor cocktail) and incubated on ice for 20 minutes, followed by sonication of the sample. Next, the sample was centrifuged at 1700g for 5 minutes at 4°C, and the supernatant was collected as the chromatin fraction. Thereafter, protein quantification was conducted utilizing a BCA reagent. Proteins were ultimately separated via SDS-PAGE, followed by a Western blot analysis employing the corresponding antibodies.

### *In silico* oligomer formation and molecular docking

The *in-silico* oligomerization was conducted utilizing the structure of the TIP60 protein (full-length). The complete structure was created via the RoseTTAFold module of the Robetta server. The produced structure was improved utilizing the GalaxyRefine tool. The structure was then verified utilizing the ERRAT, Verify3D, and PROCHECK tools accessible on the SAVES server (version 6.0) and the ProSA web server. Additionally, MD simulation was carried out to stabilize the structure using BIOVIA Discovery Studio. Finally, the GalaxyHomomer online tool was utilized to forecast the oligomeric state of full-length TIP60 protein (Baek et al., 2017). For generating mutations in the full-length TIP60, the mutagenesis tool of PyMOL was utilized. Further, to predict the oligomeric state of the mutant protein, the GalaxyHomomer server was used.

For docking studies, we utilized the CDocker module of Biovia Discovery Studio. All the default parameters (Pose Cluster Radius=0.5, Random Conformations=100, Parallel Processing=True, Processors=12) were used for performing the docking analysis. The ligands for the docking were obtained from PubChem (acetyl coenzyme A (CID – 444493), lactyl coenzyme A (CID – 3081970), crotonyl coenzyme A (CID – 5497143), propionyl coenzyme A (CID – 92753), and butyryl coenzyme A (CID – 122283)) (Kim et al., 2023).

### Cell survival assay

For the cell viability assay, 50,000 Huh7 cells were inoculated per well in a 12-well plate. Subsequent to their attachment, cells were transfected with the specified plasmid constructs utilizing Lipofectamine 2000. The medium was changed 5 hours after transfection. Following a 24-hour media replacement, cells were treated with 3µM of doxorubicin for a duration of 6 hours. Post-treatment, medium was replaced and the cells were cultured for an additional 48 hours, after which surviving cells were enumerated using a Neubauer chamber.

### Real time PCR

Huh7 cells were transfected with the specified plasmid constructs utilizing Lipofectamine 2000 according to the manufacturer’s protocol. Following a 24-hour media replacement, DNA damage was induced by treating the cells with 3µM doxorubicin for a duration of six hours and the media was then replaced and culture was maintained for an additional 24 hours. Cells were harvested after 24 hours for RNA isolation using TRIzol reagent. The isolated RNA was then treated with DNase I and subsequently quantified and utilized for cDNA synthesis using the Verso cDNA synthesis kit in accordance with the manufacturer’s guidelines. QPCR was conducted utilizing gene-specific primers, synthesized cDNAs, and SYBR green master mix with the QuantStudio 5 Real-Time PCR System (Applied Biosystems, USA). The Ct values obtained were utilized to calculate relative mRNA expression using the formula: relative quantification = 2^(-ΔΔCt). GAPDH served as an endogenous control, and gene expression was determined following normalization of values with GAPDH.

### Multiple sequence alignment and statistical analysis

Multiple sequence alignment was conducted utilizing Clustal Omega/Clustal W (https://www.ebi.ac.uk/jdispatcher/msa/clustalo). All statistical analyses were conducted utilizing GraphPad Prism 8.0 software.

## RESULTS

### Cancer-associated arginine mutations at position 53 and 62 in TIP60’s chromodomain alters its conformational stability

Since TIP60s chromodomain plays a critical role in chromatin association, we sought to determine the impact of oncogenic mutations specifically located within this domain on its functional dynamics. For this, missense mutants of TIP60 were systematically mined from cBioPortal, which aggregates large-scale cancer genomics data, and from the COSMIC database, which catalogs patient-derived somatic mutations. This approach revealed six chromodomain alterations detected in different cancers reported by cBioPortal and 22 by COSMIC database, with six recurrent mutations overlapping across both datasets, which were eventually selected for further studies (**Figure 1A, 1B**). Among the 6 identified mutations, PredictSNP analysis, which integrates multiple algorithms (MAPP, PolyPhen-1, SIFT, PhD-SNP, PolyPhen-2, SNAP), consistently identified R53H and R62W as deleterious (**Figure 1C**). Stability predictions using I-Mutant 2.0, SAAFEC-SEQ, and INPS yielded negative ΔΔG value indicating destabilization, while AlphaFold pathogenicity indexing supported their disruptive nature by classifying both variants as pathogenic in nature (**Figure 1D, 1E**).

**Figure 1:**
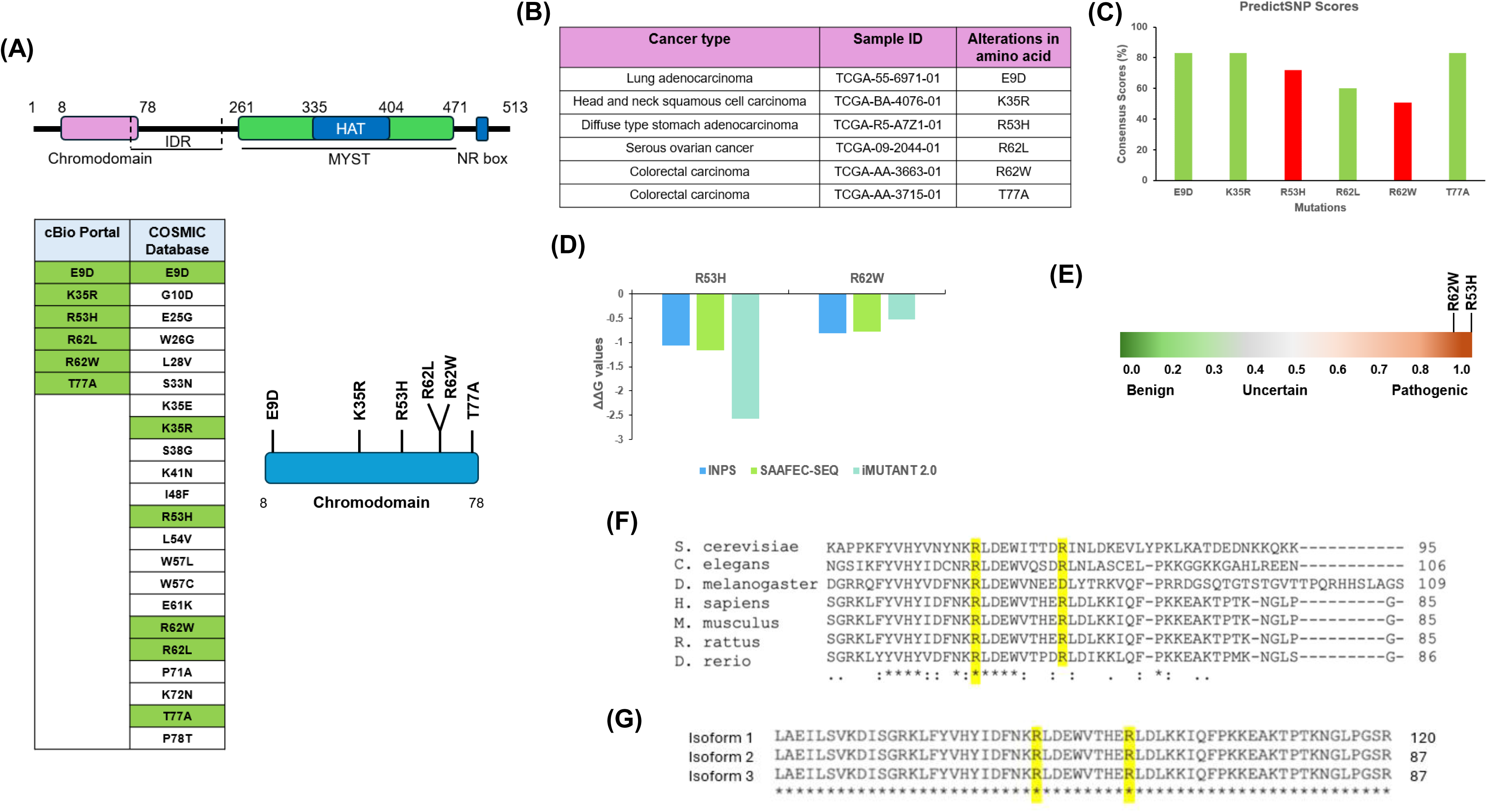
Screening and identification of TIP60’s chromodomain associated oncogenic mutations. (A) Schematic diagram of TIP60 depicting its different domains. List of cancer-associated mutations located in chromodomain from cBio Portal & COSMIC databases. Green highlights the mutations common in both the databases. (B) Table shows identified TIP60’s chromodomain mutations in different cancer. (C) Bar graph illustrating the Predict SNP consensus score (% values denote consensus scores from six distinct tools) for chromodomain mutations. Red bar represents mutations anticipated to adversely affect the functions of the TIP60 protein. (D) Bar graph illustrating the variation in Gibbs free energy (ΔΔG) of selected TIP60 mutations. All values cited are in Kcal/mol, with negative values signifying instability. (E) Heat map represents the pathogenicity index of selected TIP60 mutations. R53H has a pathogenic index of 1.0, and R62W has a pathogenic index of 0.95. (F & G) Multiple sequence alignment of the TIP60 protein across different species, highlighting the conserved arginine at position 53 and 62 (shown in yellow) and in three isoforms of TIP60 respectively.

To assess the evolutionary conservation of these oncogenic mutation sites identified in the TIP60 chromodomain, we performed multiple sequence alignment across representative eukaryotic species including *Saccharomyces cerevisiae*, *Caenorhabditis elegans*, *Drosophila melanogaster*, *Homo sapiens*, *Mus musculus*, *Rattus rattus*, and *Danio rerio*. This analysis revealed that the two residues of interest are highly conserved across all examined species, underscoring their functional importance (**Figure 1F)**. In addition, alignment of the three known human TIP60 isoforms confirmed that these positions remain invariant, thereby excluding isoform-specific variability and reinforcing the critical nature of these sites (**Figure 1G**).

To enable comparative evaluation against the wild-type TIP60 chromodomain, structural models of the TIP60 wild-type chromodomain (residues 1-78) as well as TIP60 chromodomain carrying either of the two deleterious mutations (R53H and R62W) were generated using RoseTTAFold, and the top model showed close alignment with a known crystal structure, reflected by a low RMSD, thereby validating its reliability for subsequent mutation analyses (**Figure 2A, superimposition not shown**). Comparison of the mutant structures for R53H and R62W with the wild-type, revealed minimal backbone deviations and no overt static disruption **(Figure 2B, 2C)**. Recognizing that static models cannot capture dynamic fluctuations under physiological conditions, molecular dynamics simulations of 500 ns were performed using the Desmond suite for wild-type and mutant chromodomains, followed by quantitative trajectory analyses. Root-mean-square deviation (RMSD) analysis of Cα atoms demonstrated markedly greater deviations in both mutants, indicating reduced conformational stability and enhanced structural drift (**Figure 2D**). Radius of gyration (Rg) values were consistently found to be lower in mutants, suggesting a more compact fold, which was corroborated by reduced solvent-accessible surface area (SASA) and increased burial of hydrophobic residues (**Figure 2E, 2F**). Intramolecular hydrogen bond enumeration further reinforced these observations, as both mutants formed more internal hydrogen bonds than wild-type, stabilizing the compacted state through augmented interaction networks (**Figure 2G**). Finally, residue-level root-mean-square fluctuation (RMSF) of Cα atoms highlighted elevated flexibility at multiple positions in the mutants relative to wild-type, revealing localized dynamic instability amid the globally tighter fold (**Figure 2H**). Collectively, these results indicate that R53H and R62W mutations alter the conformational dynamics of the TIP60 chromodomain, driving a shift toward a compact yet dynamically unstable fold.

**Figure 2:**
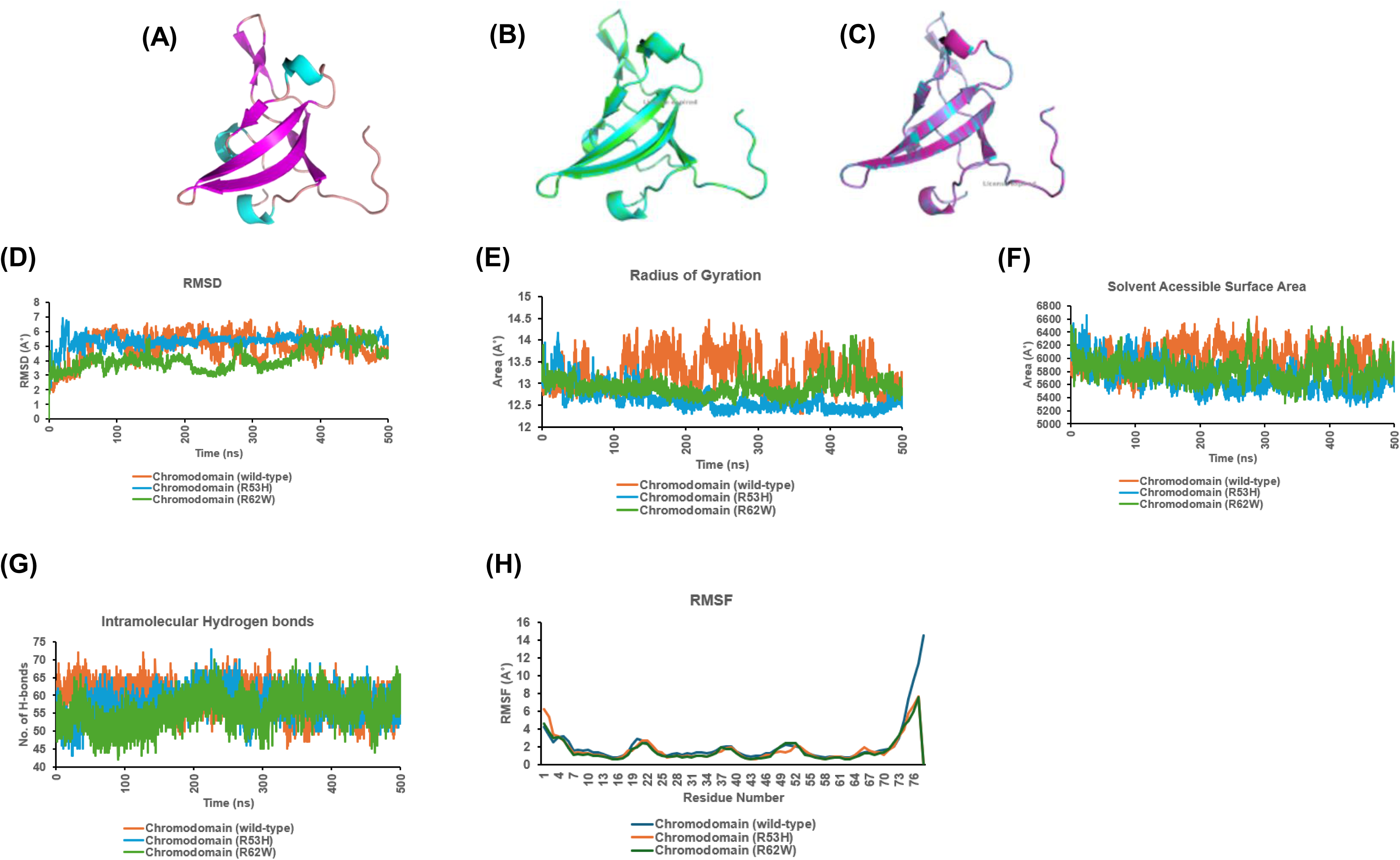
R53H and R62W affects chromodomain structure. (A) Ribbon diagram of TIP60 Chromodomain wild-type (1-78 amino acid). (B) Superimposition of modeled TIP60 (R53H) (represented in green) and TIP60 (wild-type) (represented in cyan) structures with RMSD of 0 Å. (C) Superimposition of modeled TIP60 (R62W) (represented in magenta) and TIP60 (wild-type) (represented in cyan) structures with RMSD of 0 Å. (D) RMSD graph of Cα atoms for TIP60 wild-type and mutant proteins indicating conformational stability. (E) Rg graph illustrating the TIP60 WT and mutant proteins compactness. (F) Graph of Solvent Accessible Surface Area (SASA) for wild-type and mutant proteins. (G) Graph depicts the quantity of intramolecular hydrogen bonding in relation to time for indicated proteins. (H) The RMSF (Root Mean Square Fluctuation) graph of Cα atoms for wild-type and mutant TIP60 proteins illustrating the residue wise flexibility of protein structure.

### R53H and R62W mutations do not affect TIP60’s subcellular dynamics and chromatin association

To determine whether the structural destabilization predicted for the R53H and R62W mutations manifests as altered subcellular dynamics, disrupted expression patterns, or impaired chromatin loading, we cloned these mutant variants into RFP expression vectors, designating them RFP-TIP60 (R53H) and RFP-TIP60 (R62W). Western blot analysis confirmed that both RFP-TIP60 (R53H) and RFP-TIP60 (R62W), mutants were expressed at the expected molecular weight and at levels comparable to wild-type RFP-TIP60 (**Figure 3A**). We next performed live cell imaging to determine the subcellular localization of these mutants in Cos-1 cells transiently transfected with wild-type or CD mutant constructs of TIP60. Live-cell imaging results revealed that both mutants localized to the nucleus and formed characteristic nuclear foci indistinguishable from wild-type, suggesting that these oncogenic mutations in CD does not affect the nuclear targeting and foci formation of TIP60 (**Figure 3B**). Given that these mutations reside within the chromodomain, we next wanted to probe the impact of these mutations on chromatin binding ability of TIP60 by performing subcellular fractionation assays. For this, soluble and chromatin fractions were prepared from Cos-1 cells expressing wild-type and CD mutant versions of TIP60 followed by Western blot analysis. Both mutant proteins were detected in the chromatin-bound fraction at the levels similar to the wild-type TIP60 (**Figure 3C**). GAPDH and histone H4 served as markers for soluble and chromatin-associated fractions, respectively. Taken together, these results demonstrate that despite the destabilizing effects observed *in silico*, R53H and R62W mutations do not impair TIP60 expression, nuclear localization, or chromatin binding under the tested conditions.

**Figure 3:**
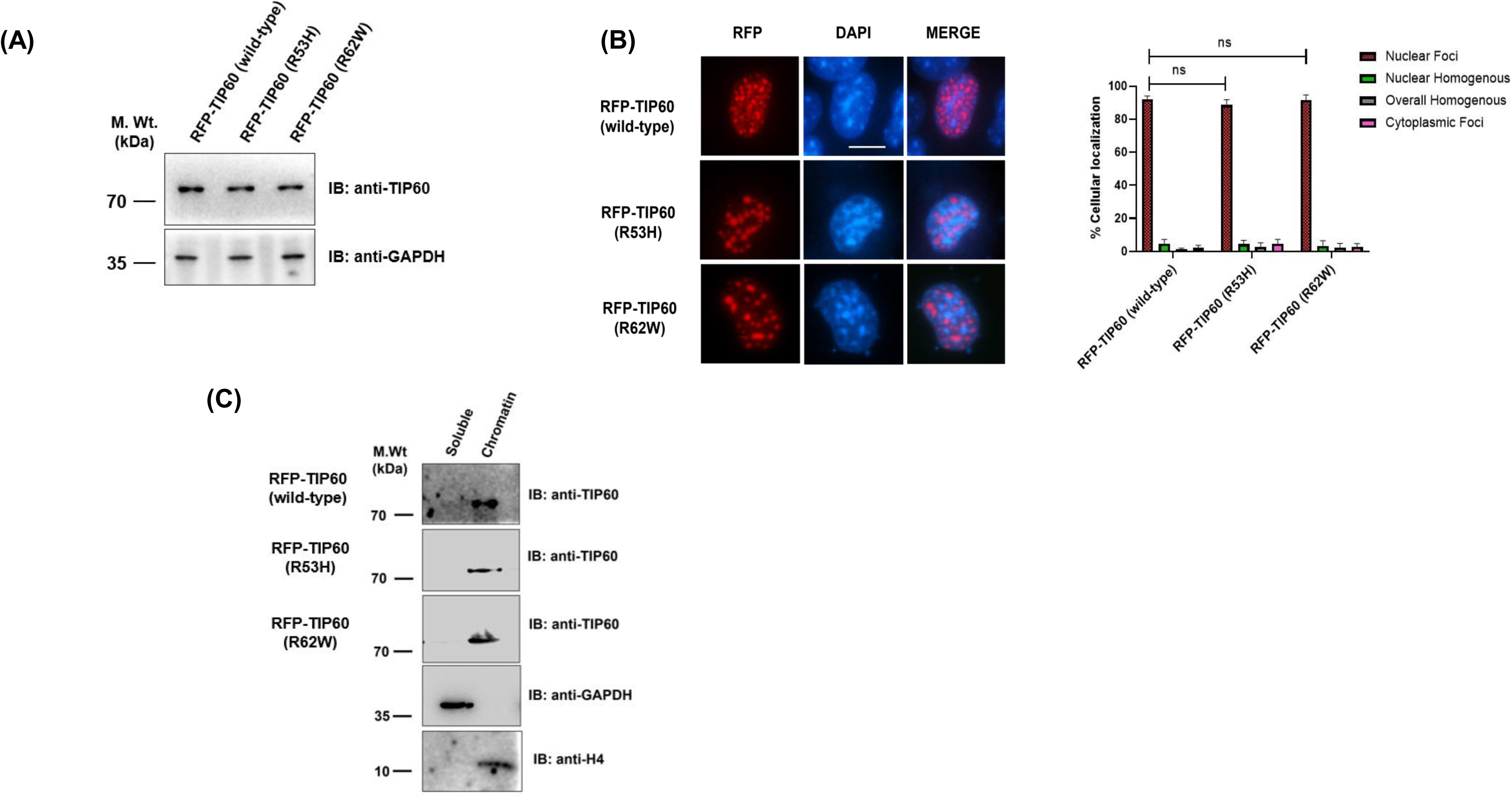
Chromodomain mutations (R53H & R62W) do not affect TIP60 intracellular localization and chromatin binding. (A) Western blot demonstrating the expression of TIP60 wild-type and mutant proteins. GAPDH served as a loading control. (B) Live cell imaging in Cos-1 cells demonstrating the intracellular distribution of RFP-tagged wild-type and mutant TIP60 proteins (red). DAPI shows nuclear staining (blue). The scale bar indicates 10μm. Graph illustrates the subcellular distribution pattern of the expressed protein within the nucleus of transfected cells. **(C)** COS-1 cells transfected with RFP-TIP60 (wild-type), RFP-TIP60 (R53H), or RFP-TIP60 (R62W) plasmids were subjected to subcellular fractionation. The isolated fractions were analyzed by Western blotting using the indicated antibodies. GAPDH and histone H4 served as fractionation controls for the soluble and chromatin fractions, respectively.

### Oncogenic mutation in TIP60’s chromodomain impairs its catalytic function

Following structural analyses, and given that nuclear localization and chromatin loading of TIP60 CD mutants remained intact, we next examined whether R53H and R62W mutations compromise TIP60’s enzymatic competence. To assess this, recombinant His-tagged TIP60’s wild-type or chromodomain mutant variants were expressed and purified from *E. coli*, and was subjected to *in vitro* histone acetyltransferase (HAT) and autoacetylation assays. For *in vitro* HAT assay, recombinant histone H4 was used as substrate, and TIP60-mediated acetylation status at lysines K5, K8, K12, and K16 was monitored by Western blot analysis. Both mutants exhibited reduced acetylation for each of these marks compared to wild-type, with R62W showing a more pronounced loss of acetyltransferase activity (**Figure 4A**).

**Figure 4.**
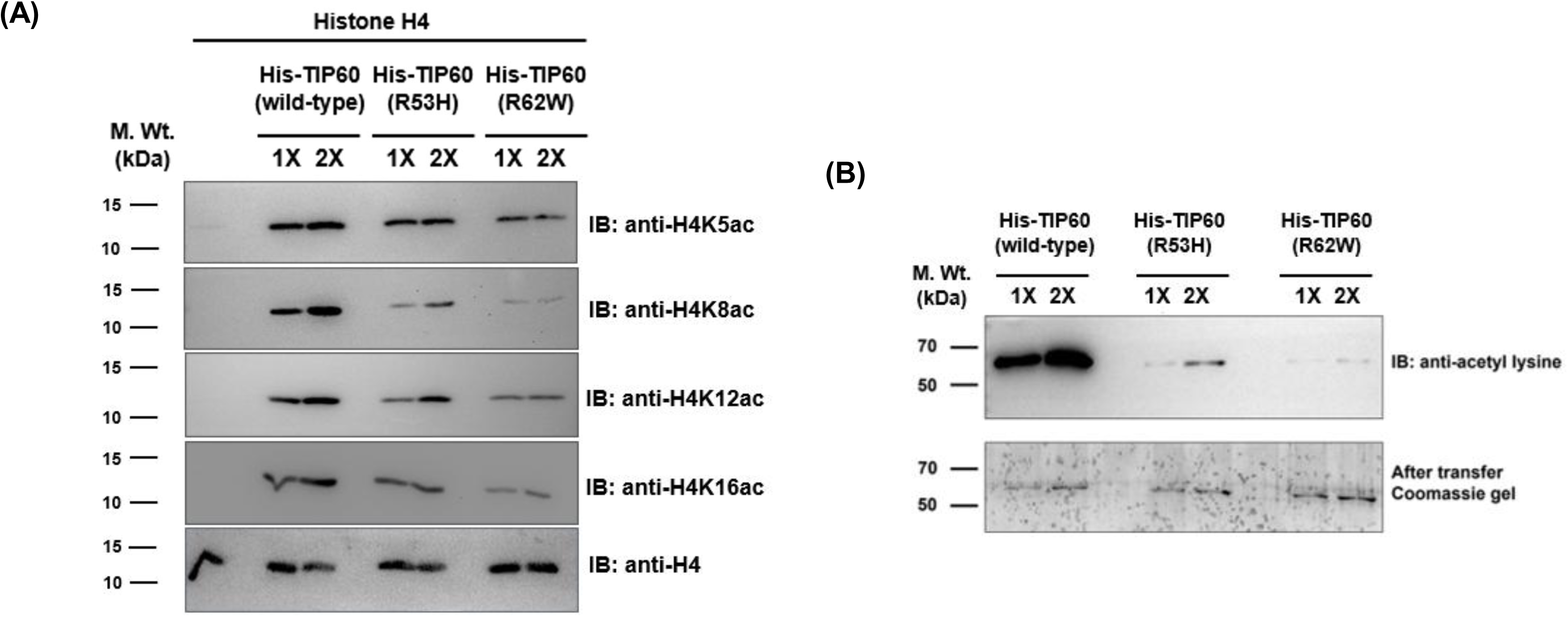
TIP60 chromodomain mutations impair its catalytic functions. (A) An *in vitro* histone acetyltransferase (HAT) assay was performed using purified recombinant His-TIP60 (wild-type), His-TIP60 (R53H), and His-TIP60 (R62W) mutant proteins in the presence of histone H4 as a substrate. Western blot analysis was carried out using antibodies against acetylated histone H4 at K5, K8, K12, and K16, along with total histone H4 as a control. (B) *In vitro* autoacetylation assays were performed using the indicated purified recombinant proteins. Western blotting was conducted using an anti-acetyl-lysine antibody. Following protein transfer, Coomassie staining of the gel confirmed equal loading of wild-type and mutant protein samples used for Western blot analysis.

Because autoacetylation is a critical self-regulatory mechanism intrinsic to TIP60, we also evaluated the TIP60’s autoacetylation activity using the same purified proteins and found that both mutants displayed lower autoacetylation levels than wild-type TIP60, consistent with a general reduction in enzymatic function (**Figure 4B**). In line with the HAT assay, the R62W mutant exhibited a greater reduction in autoacetylation signal than the R53H mutant (**Figure 4B**). Together, these results demonstrate that while R53H and R62W do not interfere with TIP60’s expression or its ability to localize to nucleus or bind chromatin, they affect its enzymatic function. This suggests that chromodomain integrity is not only essential for maintaining structural stability but also for sustaining catalytic activity, and that perturbations at conserved arginine residues, particularly R62W, translate into measurable defects in histone acetylation and self-acetylation capacity.

### R62W mutation in TIP60’s chromodomain destabilize TIP60’s trimeric oligomer formation essential for its catalytic activity

Following the observed reductions in histone acetyltransferase (HAT) and autoacetylation activities for the R53H and R62W chromodomain mutants, *in silico* modelling and docking studies were performed to understand the underlying structural basis for these enzymatic defects. TIP60 relies on binding acetyl-CoA as a cofactor to transfer acetyl groups to lysine substrates, so disruptions in this interaction could directly impair catalysis. Initial docking attempts with a full-length monomeric TIP60 model failed to produce stable acetyl-CoA binding poses (data not shown), consistent with prior evidence that TIP60 predominantly functions as a trimeric oligomer (Dubey et al., 2024; Gupta et al., 2025). Moreover, mutations can change the oligomerization capability of TIP60 (Dubey et al., 2024; Gupta et al., 2025). To investigate mutation-induced oligomerization effects, homotrimeric models of wild-type and mutant TIP60 were generated using the Galaxy server. The wild-type structure adopted a stable trimer configuration (**Figure 5A, Top panel**), with specific interfacial residues across chains, including 239, 323, 375, 466, 468, and 472 position amino acids (**Figure 5B, top panel**). Both mutants also assembled into trimers (**Figure 5A, middle and bottom panels**). The R53H trimer preserved the wild-type interfacial residue set (239, 323, 375, 466, 468, 472), showing no changes at the monomer-monomer contacts (**Figure 5B, middle panel**). The R62W trimer, however, displayed an altered profile with residues 20, 52, 166, 168, 169, 211, 253, 363, and 511 contributing to interfaces, indicating mutation-induced shifts in inter-monomer engagement (**Figure 5B, bottom panel**).

**Figure 5.**
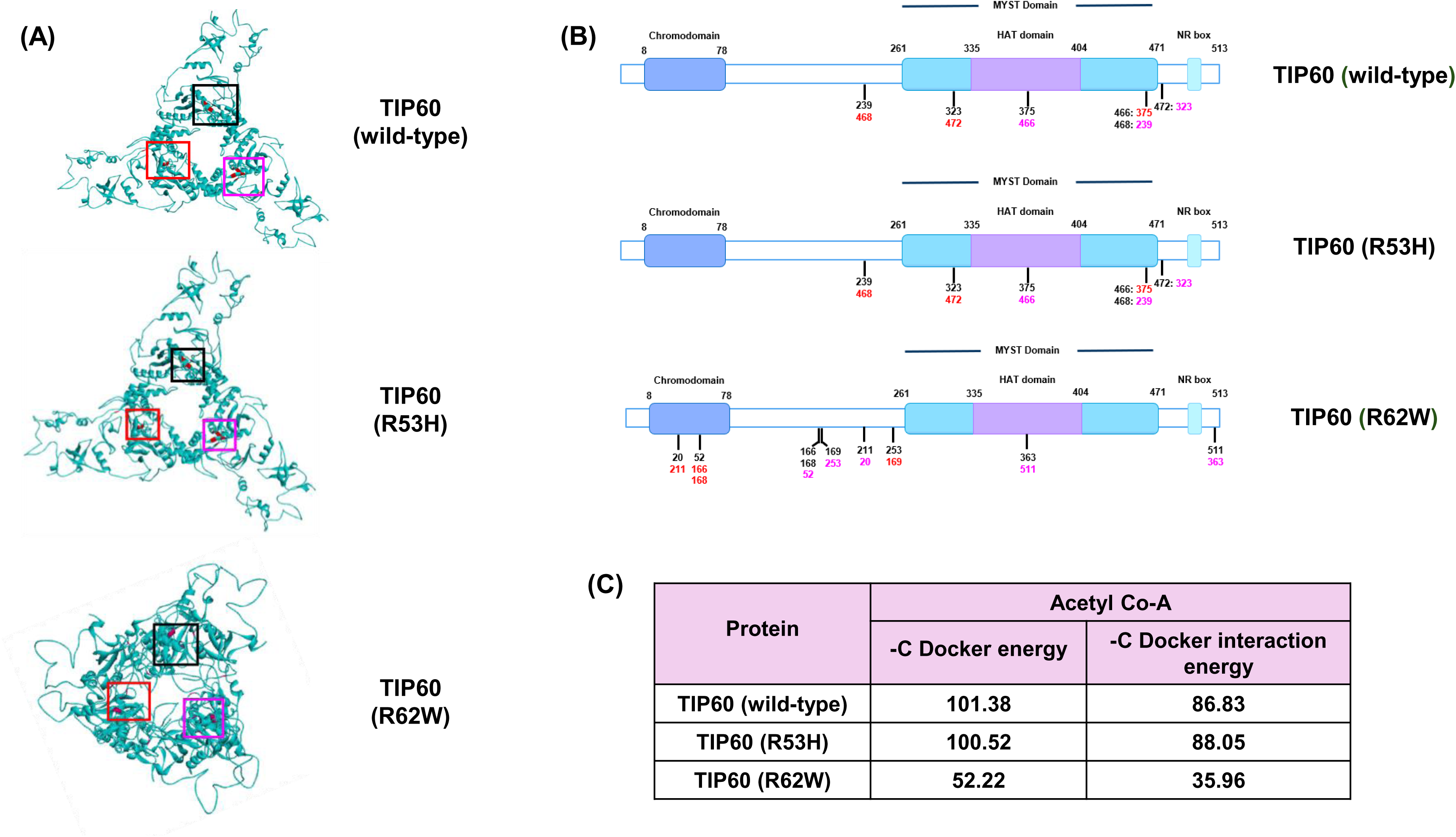
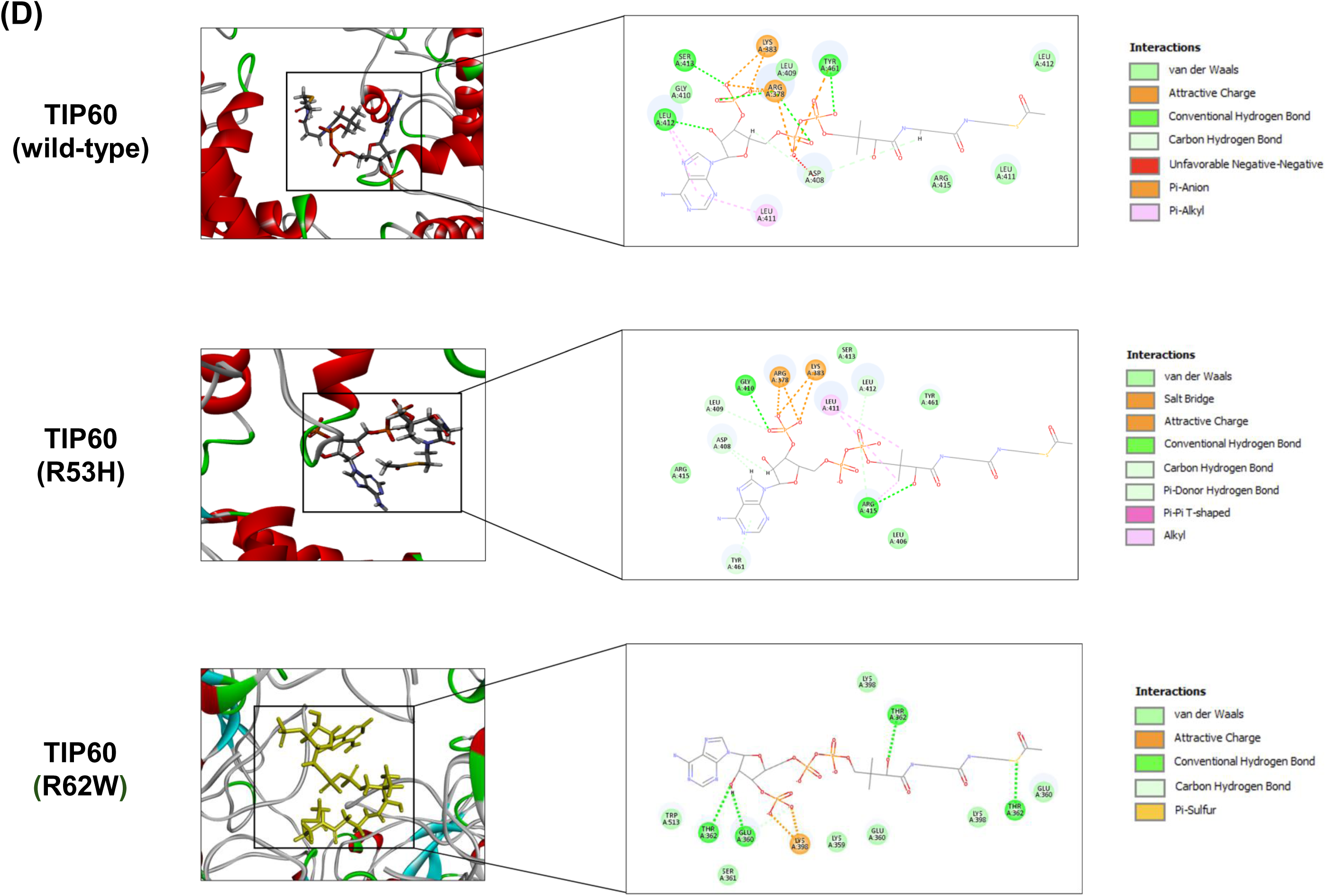
TIP60 docks with acetyl-coenzyme A moieties in a trimeric conformation. (A) Structural representation of the trimeric forms of TIP60 (wild-type), TIP60 (R53H), and TIP60 (R62W). Black, red, and magenta boxes indicate the interaction interfaces between adjacent monomeric units. (B) Linear schematic illustrating the amino acid residues involved in inter-monomer interactions within TIP60 (wild-type), TIP60 (R53H), and TIP60 (R62W) trimers, along with the specific residue-level contacts. (C) Table summarizing the binding energy values for acetyl-coenzyme A in complex with TIP60 wild-type and chromodomain mutants. More negative values indicate stronger binding affinity. (D) Structural depiction of acetyl-coenzyme A positioned within the binding pocket of trimeric TIP60. Insets show magnified views of the bonding patterns and key residues involved in interactions between the TIP60 trimer and acetyl-coenzyme A for TIP60 (wild-type), TIP60 (R53H), and TIP60 (R62W).

Trimeric models were subsequently docked with acetyl-CoA using CDocker. The wild-type trimer bound acetyl-CoA robustly, achieving a CDocker energy of -101.38 kcal/mol and interaction energy of -86.83 kcal/mol, stabilized by specific interactions including hydrogen bonds and hydrophobic contacts at the binding site (**Figure 5C**). The R53H trimer maintained productive binding, with a CDocker energy of -100.52 kcal/mol and interaction energy of -88.05 kcal/mol, reflecting only minor perturbations despite equivalent trimer interfaces (**Figure 5C**). In marked contrast, the R62W trimer exhibited severely weakened binding, with CDocker energy dropping to -52.22 kcal/mol and interaction energy to -35.96 kcal/mol (**Figure 5C**). Detailed inspection revealed disrupted hydrogen bonding networks and poorer hydrophobic enclosure at the acetyl-CoA site, attributable to the reconfigured trimer interfaces that alter pocket geometry (**Figure 5D**).

Extending this analysis, docking with alternative acyl-CoA cofactors (lactyl-CoA, crotonyl-CoA, butyryl-CoA, propionyl-CoA) succeeded only in the trimeric state, underscoring oligomerization as a prerequisite for diverse acyltransferase activities. Again, R62W exhibited reduced binding affinities across all cofactors compared to wild-type and R53H, likely due to the same interfacial perturbations that compromise pocket integrity (**Supplementary Table 3**). These findings reveal that while both mutations preserve trimer formation, the R62W mutation uniquely remodels interfacial contacts to destabilize the acetyl-CoA binding pocket, explaining its greater enzymatic deficits, while R53H preserves quaternary integrity and cofactor accommodation. Additionally, these results demonstrate that chromodomains extend beyond their role as chromatin-binding modules, exerting allosteric control over TIP60’s catalytic function by stabilizing trimeric oligomer formation.

### Chromodomain mutations abolish TIP60-mediated p21 transactivation and sensitize cells to doxorubicin

TIP60 is known to acetylate ATM and p53 during DNA damage, thereby promoting repair pathways, maintaining genomic stability, and driving *p21* transactivation through p53 activation. To determine whether chromodomain mutations compromise these responses, we first examined their impact on *p21* induction. For this, Huh-7 cells were transfected with RFP-TIP60 (wild-type, R53H, or R62W) together with GFP-p53, followed by 24-hour doxorubicin treatment to induce DNA damage. RNA was extracted using TRIzol, converted to cDNA, and analysed by qPCR for *p21* transactivation. Wild-type TIP60 significantly upregulated p21 expression under damage conditions relative to controls, whereas both R53H and R62W mutants failed to induce p21, indicating defective transcriptional activation (**Figure 6A**).

**Figure 6.**
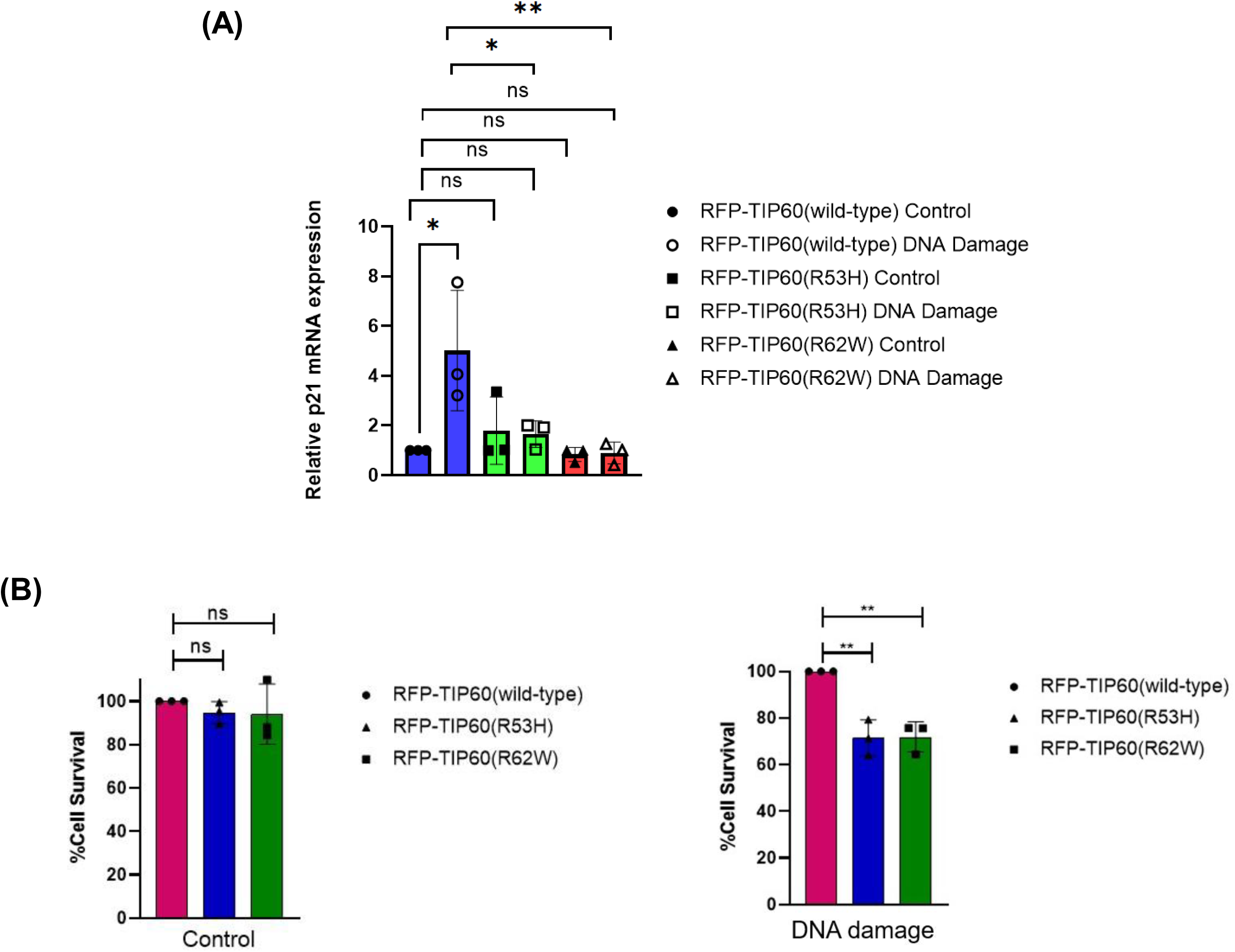
R53H and R62W mutants fail to activate p21 expression and protect cells from DNA damage. (A) RT–qPCR analysis was performed in transfected Huh7 cells to examine *p21* gene expression under DNA damage conditions compared with control. The bar graph depicts relative p21 mRNA expression, normalized to the control set (TIP60 wild-type taken as 1). GAPDH was used as an internal control. Data represent the mean ± S.D. of three independent experiments. The Y-axis indicates relative fold induction in mRNA levels. (B) Huh7 cells were transfected with the indicated plasmids. After 24 hours of protein overexpression, cells were treated with 3 μM doxorubicin for 6 hours. The medium was then replaced, and viable cells were counted after 48 hours. The left bar graph represents control (undamaged) conditions, while the right bar graph represents DNA-damaged conditions. Cell viability was calculated, and graphs were plotted using GraphPad Prism 8. Data represent mean ± S.D. of three biological replicates. ‘ns’ denotes non-significance, ‘*’ indicates p ≤ 0.05, and ‘**’ indicates p ≤ 0.01.

To assess functional consequences on DNA repair perturbations, cell viability was measured under the same conditions. For this, Huh-7 cells expressing the same TIP60 constructs with GFP-p53 were transfected and followed by treatment with doxorubicin. Cell counting data showed that in untreated controls, no differences was observed in survival rates across wild-type and mutant transfected cells (**Figure 6B, left**). However, under doxorubicin treatment condition cells expressing R53H or R62W exhibited markedly reduced viability compared to wild-type TIP60-expressing cells (**Figure 6B, Right)**. Together, these results demonstrate that chromodomain mutations abolish TIP60-mediated *p21* transactivation and fail to protect against DNA damage-induced repair, underscoring the requirement of chromodomain integrity for TIP60’s role in transcriptional activation and DNA repair pathways.

## DISCUSSION

Chromodomains are conserved modules found in diverse proteins, including HP1, Polycomb, and CHD family members, where they mediate recognition of histone modifications and facilitate chromatin association essential for transcriptional regulation and genome organization (Eissenberg, 2012). For instance, the chromodomain of HP1 helps it bind specifically to H3K9-methylated histone tails, enabling stable association with heterochromatin critical for driving transcriptional repression. Mutations within chromodomain-containing proteins have been implicated in various cancers, and other diseases often leading to defective chromatin remodelling and altered gene expression programs (Kim et al., 2008; Sharma et al., 2025). Chromodomain mutations in CHD4, identified in endometrial and colorectal cancers, impair nucleosome remodeling and DNA repair to promote tumorigenesis (Li et al., 2018; Lin et al., 2019), while CBX7 variants weaken H3K27me3 binding and trigger aberrant activation (Clermont et al., 2014; Nichol et al., 2016).

Given the pathogenic relevance of chromodomain mutations across human disease through their broad effects on protein function and chromatin interactions, we hypothesized that cancer-associated variants in the TIP60 chromodomain would impair its functional dynamics, especially its chromatin-binding ability. Contrary to this assumption, the two mutations we tested did not impair chromatin loading of TIP60, but instead, they profoundly compromised its catalytic activity, including autoacetylation and histone acetyltransferase (HAT) function. Considering that chromodomain and MYST catalytic domain are structurally separated, bridged by a flexible intrinsically disordered region (IDR), we found it remarkable that a single point mutation in the chromodomain, positioned far from the catalytic site, allosterically disrupted TIP60’s catalytic activity **(Figure 4)**. On further investigation, we found that the structural destabilization induced by R62W mutation compromised TIP60’s trimeric oligomer assembly, a prerequisite for acetyl-CoA binding and catalytic function. While the R53H mutation did not overtly affect trimerization, it nevertheless abolished TIP60’s catalytic activity. This effect likely reflects subtle conformational fluctuations, as suggested by RMSD analysis, which may cause dissociation of acetyl-CoA or inefficient catalysis. Future molecular dynamics simulations of the oligomeric TIP60 complex could provide deeper insights into how these perturbations destabilize acetyl-CoA binding. These observations point to a functional interdependence between the chromodomain and MYST catalytic domain, whereby disruption of either domain compromises TIP60’s enzymatic activity and underscores the reliance of TIP60 on coordinated communication across its modular architecture. While previous studies established that MYST domain disruption impairs TIP60 chromatin loading, our work extends this framework by showing that conserved chromodomain mutations selectively decouple chromatin engagement from catalytic activity. These further highlights the significance of interdomain communication in proper assembly of TIP60’s oligomers into trimeric complex and its proper functioning.

Functionally, despite being properly loaded onto chromatin, both chromodomain (CD) mutants failed to transactivate *p21* in response to DNA damage, consistent with a loss of catalytic activity rather than impaired chromatin association. TIP60 is known to acetylate histone H4 at lysine 16 and p53 at lysine 120, modifications essential for transcriptional activation of DNA damage response genes such as p21, which halts the cell cycle to allow repair. Loss of TIP60 catalytic activity, even in the presence of intact chromatin binding, prevents p21 induction, leaving DNA lesions unrepaired and promoting mutation accumulation.

Collectively, our findings identify a novel mechanism of TIP60 dysfunction in which chromodomain mutations allosterically compromise catalytic activity through disruption of trimerization and acetyl-CoA engagement, rather than chromatin loading. This work underscores the critical interdependence of TIP60’s functional domains and demonstrates how single mutations can destabilize protein architecture, impair enzymatic function, and compromise the DNA damage response **(Figure 7)**.

**Figure 7:**
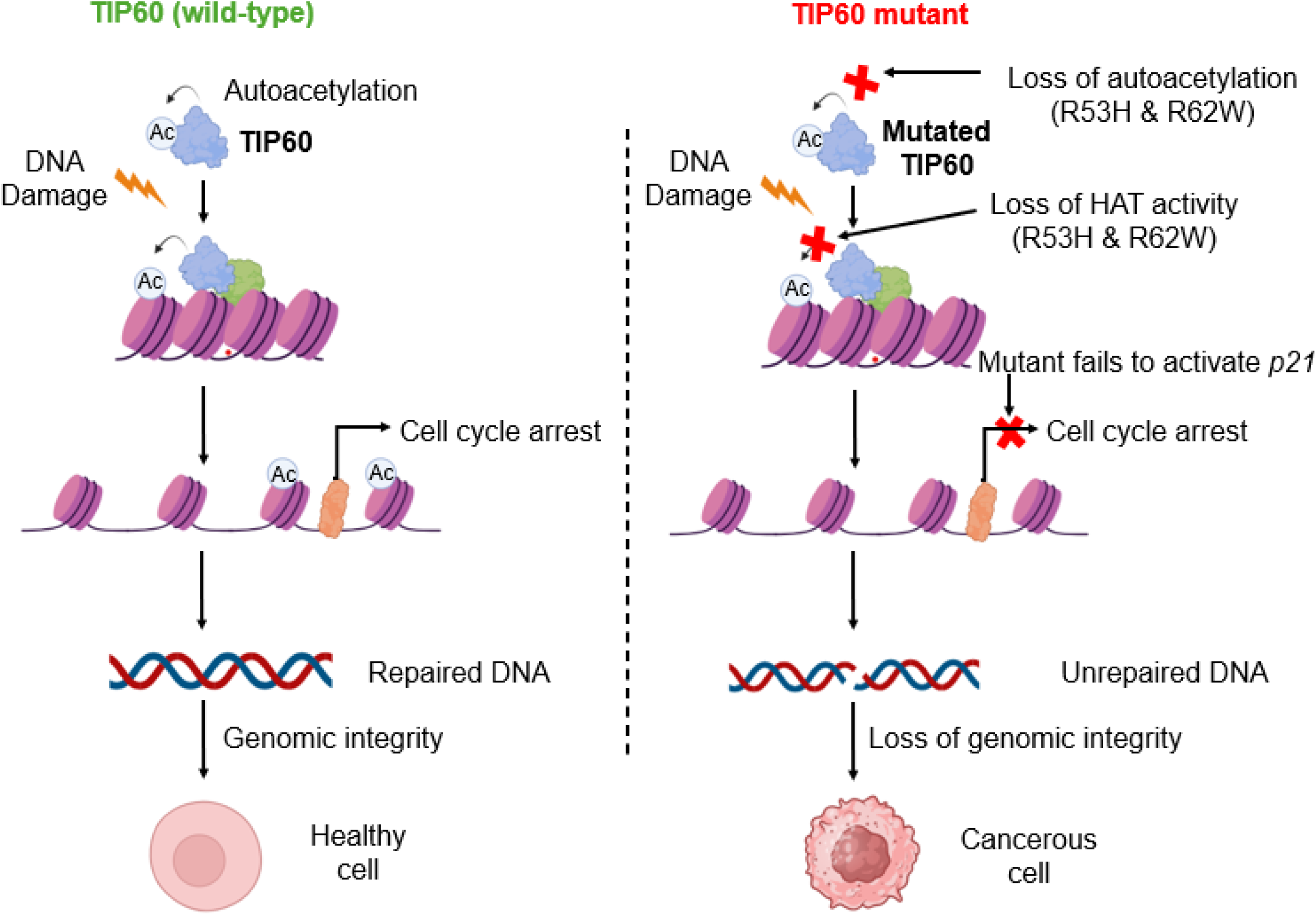
Proposed model for loss of TIP60 tumor-suppressor activity driven by chromodomain mutations. Under normal conditions, TIP60 acetylates p53, leading to the activation of p21, which promotes cell-cycle arrest and facilitates DNA repair, thereby preserving genomic integrity. In contrast, oncogenic chromodomain mutations in TIP60 (R53H and R62W) are defective in transactivating p21 under DNA-damage conditions, compromising cell-cycle checkpoint control and the DNA repair capacity of the cell. This dysfunction jeopardizes genomic integrity and may contribute to cancer initiation and progression.

By delineating this mechanism, our study expands the conceptual framework for understanding TIP60 dysfunction in cancer pathogenesis.

## Abbreviations

TIP60: tat interactive protein 60
CD: chromodomain
HAT: histone acetyl transferase

## Acknowledgements

Authors thank SNIoE for the infrastructure and resources. HG acknowledges ICMR for Senior Research Fellowship. AB thank SNIoE for OUR (opportunities for undergraduate research) program.

## Author contributions

HG- performed the experiments for the study, data acquisition and analysis, original draft writing and editing; AB- performed the experiments for the study, data acquisition and analysis; AG- Conceived, designed, and supervised the study, funding acquisition, original draft writing, review, and editing. All authors read and approved the final version of the manuscript.

