## Supplementary Material for "Oncogenic chromodomain mutations allosterically impair TIP60 acetyltransfearse function preventing activation of DNA repair genes under genotoxic stress"

**Supplementary information**

**Figure legends**

**Supplementary Figure 1**: **Cloning of TIP60 chromodomain mutants (R53H and R62W) proteins in pDS-Red vector.** (A) Agarose gel image showing the PCR amplified TIP60 mutant ORF and the double restriction digestion of obtained clones with KpnI and BamHI demonstrates the positive clones of TIP60 (R53H) mutant. (B) Agarose gel image showing the PCR amplified TIP60 mutant ORF and the double restriction digestion of obtained clones with KpnI and BamHI demonstrates the positive clones of TIP60 (R62W) mutant.

**Supplementary Figure 2**: **Cloning, expression and protein purification of TIP60 mutants in pET-28a vector.** (A) Agarose gel image showing double restriction digestion of obtained plasmids with BamHI and EcoRI to demonstrate positive clones of pET-28a TIP60 (R53H) mutant. The dotted line in the agarose panel was used to show the image stitching. (B) Agarose gel image showing double restriction digestion of obtained plasmids with BamHI and EcoRI to demonstrate positive clones of pET-28a TIP60 (R62W) mutant. Affinity purification analysis of His-TIP60 (C) Western blot with anti-His antibody shows the expression of TIP60 mutants at the expected size. (D) Coomassie gel illustrating the protein purification profile of His-TIP60 (R53H). (E) Coomassie gel illustrating the protein purification profile of His-TIP60 (R62W). Arrow indicates the purified protein.

**Supplementary Figure 1**: **Cloning of TIP60 chromodomain mutants (R53H and R62W) proteins in pDS-Red vector.**

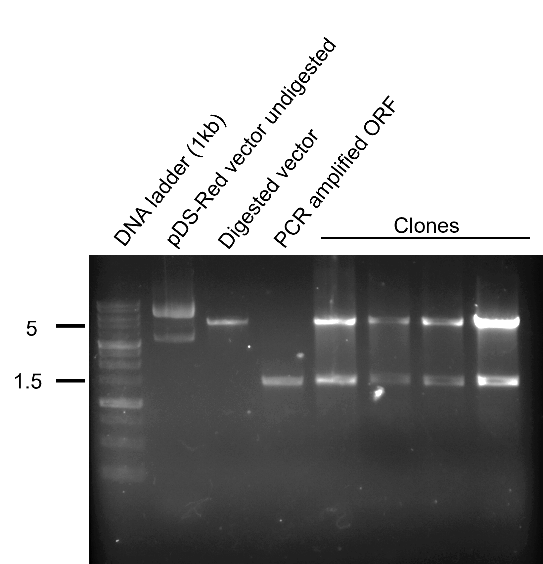

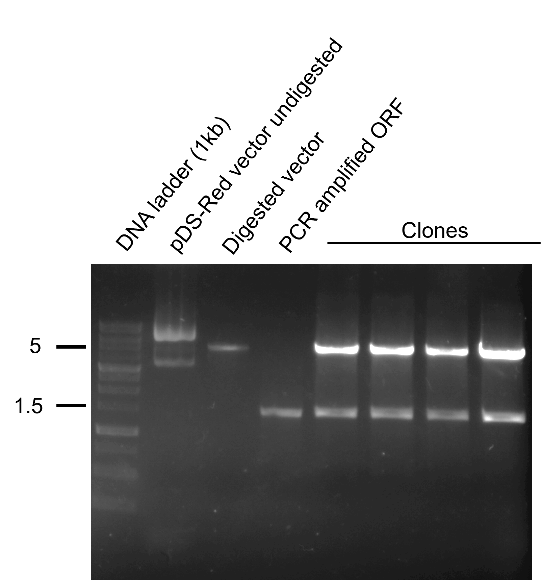

**(A)**

TIP60 (R62W)

**(B)**

TIP60 (R62W)

TIP60 (R53H)

**Supplementary Figure 2**: **Cloning, expression and protein purification of TIP60 mutants in pET-28a vector.**

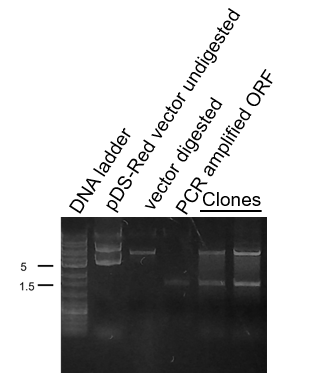

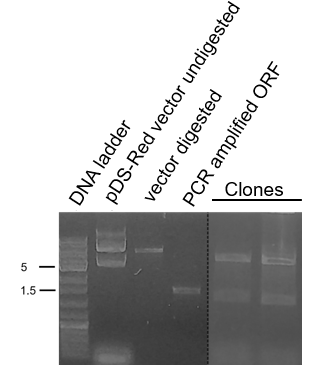

**(B)**

**(A)**

TIP60 (R62W)

TIP60 (R53H)

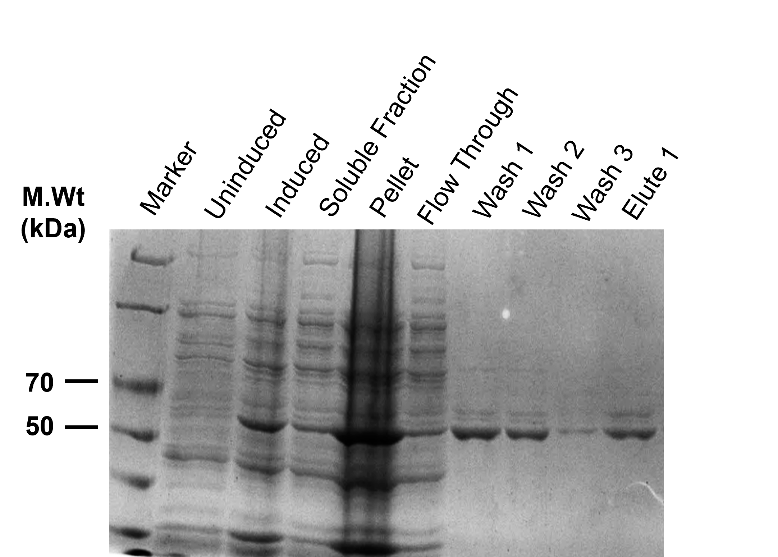

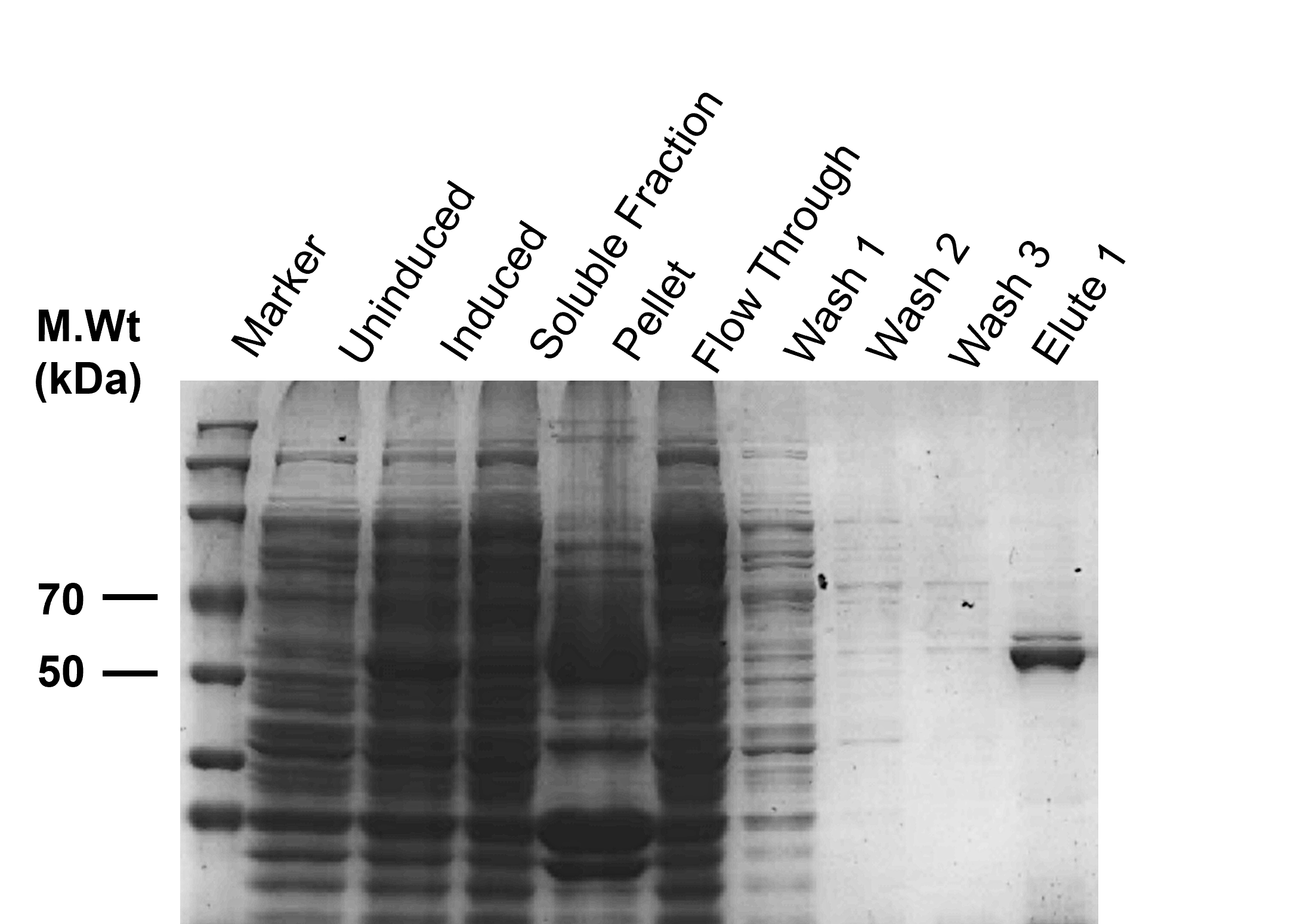

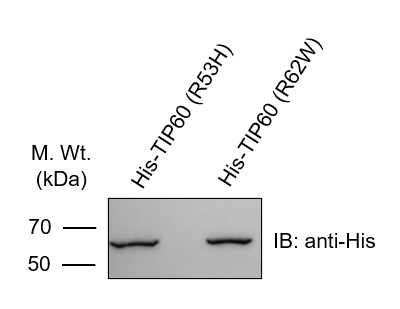

**(E)**

**(D)**

**(C)**

**Table legends**

**Supplementary Table 1: List of primers used in the study**

| **S.**  **No.** | **Primer** | **Sequence (5’-3’)** |
| --- | --- | --- |
| 1 | hTIP60-KpnI-Fw | GGGGTACCATGGCGGAGGTGGGGGAG |
| 2 | hTIP60-BamHI-Rv | CGGGATCCTCACCACTTCCCCCTCTTGC |
| 3 | hTIP60-BamHI-Fw | CGGGATCCATGGCGGAGGTGGGGGAG |
| 4 | hTIP60-EcoRI-Rv | CGGAATTCCCACTTCCCCCTCTTGCT |
| 5 | hTIP60-R53H-Fw | GACTTCAACAAACATCTGGATGAA |
| 6 | hTIP60-R53H-Rv | CCATTCATCCAGATGTTTGTTGAA |
| 7 | hTIP60-R62W-Fw | GTGACGCATGAGTGGCTGGACCTA |
| 8 | hTIP60-R62W-Rv | CTTTAGGTCCAGCCACTCATGCGT |

**Supplementary Table 2: List of antibodies used in the study.**

| **Antibodies** | **Manufacturer** | **Catalogue Number** |
| --- | --- | --- |
| 6x-His Tag Polyclonal antibody | Invitrogen | #PA1-983B |
| Acetyl-Histone H4 (K5) (D12B3) rabbit mAb | Cell Signaling Technology | #8647T |
| Acetyl-Histone H4 (Lys12) (D2W6O) rabbit mAb | Cell Signaling Technology | #13944 |
| Acetyl-histone H4 (Lys16) antibody | Affinity Biosciences | AF3636 |
| Acetyl-histone H4 (Lys8) antibody | Cell Signaling Technology | #2594 |
| Ac-lysine (AKL5C1) | Santa Cruz Biotechnology | sc-32268 |
| GAPDH (Polyclonal antibody) | Abgenex, India | NA |
| Histone H4 (D2X4V) rabbit mAb | Cell Signalling Technology | #13919S |
| m-IgGk BP-HRP | Santa Cruz Biotechnology | sc-516102 |
| Mouse anti-rabbit IgG-HRP | Santa Cruz Biotechnology | sc-2357 |
| TIP60 (C-7) | Santa Cruz Biotechnology | sc-166323 |

**Supplementary Table 3: Table mentioning the binding energy values of the acyl-coenzyme A cofactors (lactyl-CoA, crotonyl-CoA, butyryl-CoA, propionyl-CoA) along with the TIP60 wild type and mutants. Negative sign indicates more binding energy.**

| **Protein** | **Crotonyl-CoA** | | **Butyryl-CoA** | | **Lactyl-CoA** | | **Propionyl-CoA** | |
| --- | --- | --- | --- | --- | --- | --- | --- | --- |
|  | **-C Docker energy** | **-C Docker interaction energy** | **-C Docker energy** | **-C Docker interaction energy** | **-C Docker energy** | **-C Docker interaction energy** | **-C Docker energy** | **-C Docker interaction energy** |
| **TIP60 (wild-type)** | 90.946 | 86.122 | 95.48 | 87.02 | 95.76 | 87.03 | 95.65 | 98.6 |
| **TIP60 (R53H)** | 89.753 | 90.95 | 104.1 | 94.85 | 98.7 | 90.17 | 100.5 | 89.19 |
| **TIP60 (R62W)** | 47.17 | 41.914 | 54.46 | 36.27 | 59.8 | 43.8 | 60.24 | 47.91 |
